# The cyclic nucleotide binding sites of Swiss-Cheese, the *Drosophila* orthologue of human PNPLA6, are required for its catalytic function

**DOI:** 10.64898/2025.12.10.693492

**Authors:** Alexander Law, Jill Wentzell, Alexandre Bettencourt da Cruz, Luke Marney, Doris Kretzschmar

## Abstract

Mutations in Swiss-cheese (SWS) or its vertebrate ortholog PNLPA6, also called Neuropathy Target Esterase (NTE), cause progressive neuronal degeneration in *Drosophila* and mice and several complex syndromes in humans. These include mental retardation, spastic paraplegia, ataxia and blindness and several other symptoms. SWS and PNPLA6 are widely expressed in neurons and in several glial cell types in *Drosophila* and mice and both cell types require SWS/PNPLA6 function autonomously. SWS and PNPLA6 are structurally and functionally conserved because expression of human or mouse PNPLA6 can replace SWS in flies. These orthologues share several domains, including the highly conserved phospholipase domain that mediates its function in deacetylating phosphatidylcholine (PC) to lysophosphatidylcholine and glycerophosphocholine. In addition, they share three cyclic nucleotide binding sites and although about 10% of the known disease-causing mutations occur in these sites, their function is still unknown. We therefore generated mutations in these sites in SWS to address what consequences this has for the function of the protein. Mutating only one site (SWS^G558E^) results in a partially functional protein that rescues the sws knockdown and that decreases PC when overexpressed. However, mutating all three sites (SWS^ΔCNB^) renders SWS non-functional and results in an increase of PC when overexpressed, suggesting that cyclic nucleotide binding can regulate the phospholipase function.

## 1. Introduction

Swiss-Cheese (SWS) is the *Drosophila* homolog of patatin-like phospholipase domain-containing protein 6 (PNPLA6), also called Neuropathy Target Esterase (NTE) (Lush *et al*., 1998). While NTE has initially been identified as the target of organophosphate poisoning that was leading to neuropathy (Johnson, 1969), mutations in PNPLA6/NTE have now been shown to cause a variety of rare diseases, including Boucher-Neuhäuser, Gordon-Holmes, Laurence-Moon, and Oliver McFarlane Syndrome as well as an inherited Spastic Paraplegia (Kretzschmar, 2022; Nanetti *et al*., 2022; Liu & Hufnagel, 2023). Patients with these diseases show variable combinations of spasticity, cerebellar ataxia, hypogonadism, chorioretinal dystrophy, blindness and less frequently peripheral neuropathy and impaired cognitive functions (Deik et al. 2014, Kmoch et al. 2015, Synofzik et al. 2014, Topaloglu et al. 2014, (Rainier *et al*., 2008; Liu *et al*., 2024).

Mice with a brain-specific knock-out of PNPLA6 developed normally but showed neuronal degeneration and problems in motor coordination with age (Akassoglou et al. 2004). Knocking out PNPLA6 specifically in glia caused an incomplete ensheathment of Remak fibers by non-myelinating Schwann cells in the adult sciatic nerves (McFerrin *et al*., 2017). In contrast, complete loss of PNPLA6 resulted in embryonic lethality of the knockout mice around day 9 postcoitum due to failed placental development and defects in the vascular system of the yolk sac (Moser et al. 2004).

*Drosophila* lacking SWS due to a nonsense mutation, generating a protein of roughly a quarter of its original size that is not detectable in Western blotsshow neuronal and glial degeneration (Kretzschmar *et al*., 1997). This phenotype is first detectable after about a week of adulthood and progresses with further aging and the flies die prematurely. Similarly, a neuronal knockdown resulted in neurodegeneration and reduced lifespan in addition to locomotor and memory deficits (Melentev *et al*., 2021). Glia-specific knockdowns resulted in morphological changes in glial ensheathment in the central and peripheral nervous system, accompanied by axonal damage and locomotion deficits (Dutta *et al*., 2016; Ryabova *et al*., 2021). Together with the finding that expression of mouse or human PNPLA6 can restore SWS function in both cell types in *Drosophila* (Muhlig-Versen *et al*., 2005; Topaloglu *et al*., 2014; Dutta *et al*., 2016; Sunderhaus *et al*., 2019), this demonstrates a functional conservation of the fly and mammalian proteins.

Mammalian PNPLA6 and SWS contain a highly conserved domain that is required for the phospholipase function (Glynn, 2000; Moser *et al*., 2000; Quistad *et al*., 2003; Muhlig-Versen *et al*., 2005; Glynn, 2013). PNPLA6 preferably hydrolyzes phosphatidylcholine (PC) and lysophosphatidylcholine (LPC) to glycerophosphatidylcholine (Lush et al. 1998, van Tienhoven et al. 2002, Quistad et al. 2003) and *sws* mutants have increased levels of PC and LPC (Muhlig-Versen et al. 2005, Kmoch et al. 2015). SWS has also been shown to contain a domain that shows homology to the regulatory subunit of Protein Kinase A (PKA) that mediates the binding to the catalytic subunit (Bettencourt da Cruz *et al*., 2008). SWS specifically binds to the C3 catalytic domain of PKA (PKA-C3), and so does mouse PNPLA6, acting as a non-canonical regulatory subunit that inhibits PKA-C3 activity (Bettencourt da Cruz *et al*., 2008; Wentzell *et al*., 2014). Similar to canonical PKA regulatory subunits, SWS contains several predicted cyclic nucleotide binding (CNB) sites that are also conserved in the mammalian homologs (Glynn, 2000; Moser *et al*., 2000; Quistad *et al*., 2003; Muhlig-Versen *et al*., 2005; Glynn, 2013). Disease causing missense mutations can be found in both, the CNB domains and in the phospholipase domain (Synofzik *et al*., 2014; Kmoch *et al*., 2015; Liu *et al*., 2024). Supporting a functional role of the CNB are findings that expression of human PNPLA with mutations in the CNB region can only partially rescue the behavioral and degenerative phenotypes in the *sws* mutant and they cannot restore wildtype PC or LPC levels (Sunderhaus *et al*., 2019). To obtain more insights into the function of the CNB domains we have generated constructs with mutations in critical residues of the CNB domains and determined the functional consequences of expressing these constructs.

## 2. Materials and Methods

### 2.1. Drosophila stocks

The UAS-*sws*RNAi line was obtained from the VDRC stock center (w^1118^; P{GD3277}v5469) and *elav*-GAL4 from the BDSC stock center (P{GawB}elav^C155^, #458). The UAS-SWS line is described in (Muhlig-Versen *et al*., 2005; Bettencourt da Cruz *et al*., 2008). *Gli*-GAL4 was kindly provided by V. Auld (University of British Columbia, Vancouver, Canada). UAS-PKA-C3-Myc was generated by adding a myc-tag to our UAS-PKA-C3 construct described in (Bettencourt da Cruz *et al*., 2008). UAS-sws^G558E^ and UAS-sws^ΔCNB^, which contains mutations in all three CNB sites, were generated using the QuickChange Multi Site-directed mutagenesis kit (Stratagene, California) and the following primer pairs for CNB1: TTGAAGACGGTCAGGAAGGAGGAGTCGGTGACAT/ATGTCACCGACTCCTCCTTCCTGACGTCTTCAA, CNB2: GCATCCCGGCGAAATCGTTGAAGGACTGGCCATGC/GCATGGCCAGT CCTTCAACGATTTCGCCGGGATGC and CNB3: GCAAGGGGGATCTTGTGGAGATCGTTGAAATGATCACG/CGTGATCATTTCAACGATCTCCACAAGATCCCCCTTGC and ACGGAAACG TCGTGGACCACGACGGTGATGG/CCATCACCGTCGTGGTCCACGACGTTTCCGT. Flies were maintained and aged at 24°C with a 12:12h light dark cycle. Expression of the sws^G558E^ and sws^ΔCNB^ was confirmed by Western blot (Fig. S1).

### 2.2. Fast Phototaxis

Newly eclosed flies were collected daily and aged with fresh vials provided every 7 days. Males and females were aged together but tested and analyzed separately. Fast phototaxis assays were conducted in the dark as previously described in Dutta et al. (Dutta *et al*., 2016) using the countercurrent apparatus described by Benzer (Benzer, 1967) and a single light source. A detailed description of the experimental conditions can be found in Strauss and Heisenberg (Strauss & Heisenberg, 1993). Briefly, flies were transferred to the apparatus in groups of 10–15 flies, shaken to the bottom of the vial and allowed to transition toward the light in 5 consecutive runs, each lasting 6 s. Flies were then scored based on the tube they were contained in at the end of the final run. Statistical analyses were performed using GraphPad Prism (v.5 for windows, San Diego CA, USA) and One-way ANOVA with a Dunnett’s multiple comparison test to compare each experimental group to a control.

### 2.2. Neurodegeneration

To quantify vacuoles, paraffin sections for light microscopy were prepared and analyzed as described in (Sunderhaus & Kretzschmar, 2016). Briefly, whole flies were fixed in Carnoy’s solution and dehydrated in an ethanol series followed by incubation in methyl benzoate before embedding in paraffin. Sections were cut at 7μm and analyzed with a Zeiss Axioscope 2 microscope using the auto-fluorescence caused by the dispersed eye pigment. To quantify the vacuolization, we photographed the paraffin section that showed the most severe phenotype without knowing the genotype for double-blind analysis. The area of the vacuoles was then calculated in ImageJ as total pixel number and the genotype determined. Statistics were performed using GraphPad Prism 5 and One-way ANOVA with a Dunnett’s post-test comparing experimental groups to the control.

### 2.3. Western blots

For the PKA-C3 localization studies, lysates were prepared from approximately 300 heads from *elav*-Gal4>UAS-PKA-C3-Myc and incubated with 0.9mM cAMP, cGMP, or both for 60 min at 4^°^C in the presence of 5 μM protein kinase inhibitor to prevent re-association of PKA-C3 with regulatory subunits. Membrane and cytosolic fractions were then prepared following the protocol of (Orgad *et al*., 1987). Protein amounts were determined by Bradford assays (Bradford, 1976), and 12 μg was loaded per lane. Western blots were performed as described in (Tschäpe *et al*., 2002). Rabbit anti-Myc was used at 1:10,000 (kindly provided by M. Forte, OHSU).

To detect sws, 14 adult fly heads from males aged to 7 days were dissected on an ice-cold plate, homogenized in 40 μl tissue homogenization buffer (50mM tris, 250mM sucrose, 150mM NaCl, 5mM EDTA, 5mM EGTA, 1% DOC, 1% triton) with protease and phosphatase inhibitors (Cell Signaling Technology 5872S), and pelleted by centrifugation at 10,000 × g for 10 min at 4 °C. Supernatant was mixed with LDS Sample Buffer (ThermoFisher B0008) supplemented with 50 mM tris(2-carboxyethyl)phosphine (TCEP) as a reducing agent. Samples were heated to 95°C for 5 minutes and the equivalent of about three heads electrophoresed through 8% bis-tris gels (ThermoFisher NW0082) to separate proteins, which were then transferred to PVDF membranes (Millipore ISEQ85R). Membranes were blocked for 30 min with 10% nonfat dry milk dissolved in 1X TBST (Tris-buffered saline + 0.1% TWEEN-20) then probed with primary and secondary antibodies using standard western blotting procedures. Rabbit anti-sws (1:500; (Muhlig-Versen *et al*., 2005)), mouse anti-GAPDH G-9 (1:1,000; Santa Cruz sc-365062), goat anti-rabbit peroxidase conjugate (1:20,000; Jackson ImmunoResearch 111-035-144), goat anti-mouse peroxidase conjugate (1:20,000; Jackson ImmunoResearch 115-035-166). Enhanced chemiluminescent substrate (Michigan Diagnostics FWPD02) was used to visualize bands.

### 2.4. Lipid measurements

Thirty-five *Drosophila* heads from males aged to 30d were collected and pulverized in 700 µL cold dichloromethane:isopropanol:methanol at a ratio of 25:10:65 (containing 1 µg/mL EquiSPLASH LIPIDOMIX as internal standard, Avanti Research) using a ceramic bead beater for 1 min, centrifuged at 13,000 rpm for 10 min, and the top layer was collected for liquid chromatography ion-mobility time-of-flight mass spectrometry (LC-IM-TOFMS) analysis. Chromatographic separation was achieved by a Waters I-class Acquity UPLC equipped with a Waters Acquity UPLC CSH C18 column (100 × 2.1 mm, 1.7 µm). Gradient elution was performed similar to previous investigations (Cajka & Fiehn, 2017; Seufert *et al*., 2023). Briefly, mobile phase compositions were, Solvent A: acetonitrile:water =6:4 + 10 mM ammonium formate (0.631 g/L) + 0.1% formic acid and solvent B: Isopropanol:acetonitrile: water=9:1 + 10 mM ammonium formate (0.631 g/L) + 0.1 % formic acid. Negative ion mode mobile phases were, Solvent A: acetonitrile:water =6:4 + 10 mM ammonium acetate (0.77 g/L), Solvent B: Isopropanol:acetonitrile: water=9:1 + 10 mM ammonium acetate (0.77 g/L). The gradient started with 15% B and was held for 0.3 min at 15% B, followed by a 2-min linear gradient from 15% to 30 % B. The gradient was increased linearly to 50% B at 2.2 min, again increased to 80% B to 9 min, and 100% B to 9.3 min. The gradient was held to 11.5 min at 100% B and, finally, stepped back to 5% B to 11.8 min and equilibrate the column untill 14 min. Mass spectrometry was performed on a Bruker timsTOF mass spectrometer operating with an electrospray ionization source at a capillary voltage of 4,500 V, end plate offset of 500 V, nebulizer pressure of 2.2 bar, and a dry gas flow of 10 L/min at 220 ^°^C. Data was acquired with Parallel Accumulation Serial Fragmentation - Data-Dependent Acquisition (PASEF-DDA), detecting m/z 100-1350 and mobility (1/Ko) range 0.55 – 1.90 Vs/cm^2^ over a ramp time of 300 ms.(Cajka & Fiehn, 2017)

Samples were analyzed with a 3 µL injection, in triplicate, and in positive and negative ion modes. Acquired data were annotated with reference databases and the Metaboscape lipid annotator (Ross *et al*., 2023). Briefly, measured lipid features, such as precursor mass, retention time, collision cross section, and fragmentation spectra are compared to reference values in a database, assigning lipid annotations based on the degree of similarity between the experimental data and the library entries (Köfeler *et al*., 2021). The reproducibility of measurements was determined by internal standard components and shown in Fig. S2, S3. Relative quantification of lipids measured is achieved by normalization to internal standard components.

## 3. Results

### 3.1. Mutating the CNB sites decreases the ability to rescue the phototaxis deficits and neurodegeneration in neuronal sws knockdown flies

As described above, SWS and its mammalian homologs PNPLA6/NTE share several conserved domains. This includes an N-terminal transmembrane domain and the phospholipase domain that mediates the catalytic activity against PC and LPC (TM and PL in Fig. 1A). In addition, they share three putative CNB sites, with a single site near the N-terminus and two more localized next to each other near the middle of the proteins, which show homology to the regulatory subunit of PKA (Fig. 1A). To test whether the CNBs are required for the biological function of SWS, we generated two *sws* constructs with mutations in these sites. The binding sites for cAMP as well as cGMP contain conserved glycines and it has been shown that changing the glycine to glutamic acid disrupts the binding (Taylor & Kornev, 2011; Lorenz *et al*., 2017; Kim & Sharma, 2021). We therefore generated a SWS construct in which the glycines in all three CNB sites were mutated to glutamic acid (Fig. 1B, highlighted) to interfere with the binding of both cyclic nucleotides. In addition to the glycine, the cyclic nucleotide binding sites in the PKA regulatory subunit also contain a conserved arginine ten amino acids downstream of the glycine. While this arginine is not present in CNB1 and CNB2 it is present in CNB3 in SWS and its mammalian homologs and we therefore also mutated this residue to affect the cyclic nucleotide binding more efficiently (R to W, Fig. 1B). To test whether mutating a single CNB site is sufficient to affect SWS function, we generated a *sws* construct with the glycine only mutated in CNB2. CNB2 was chosen because it had the highest score of conservation when compared to the canonical *Drosophila* PKA-RII regulatory subunit (25 versus 24 for CNB3 and 13 for CNB1).

**Fig. 1.**
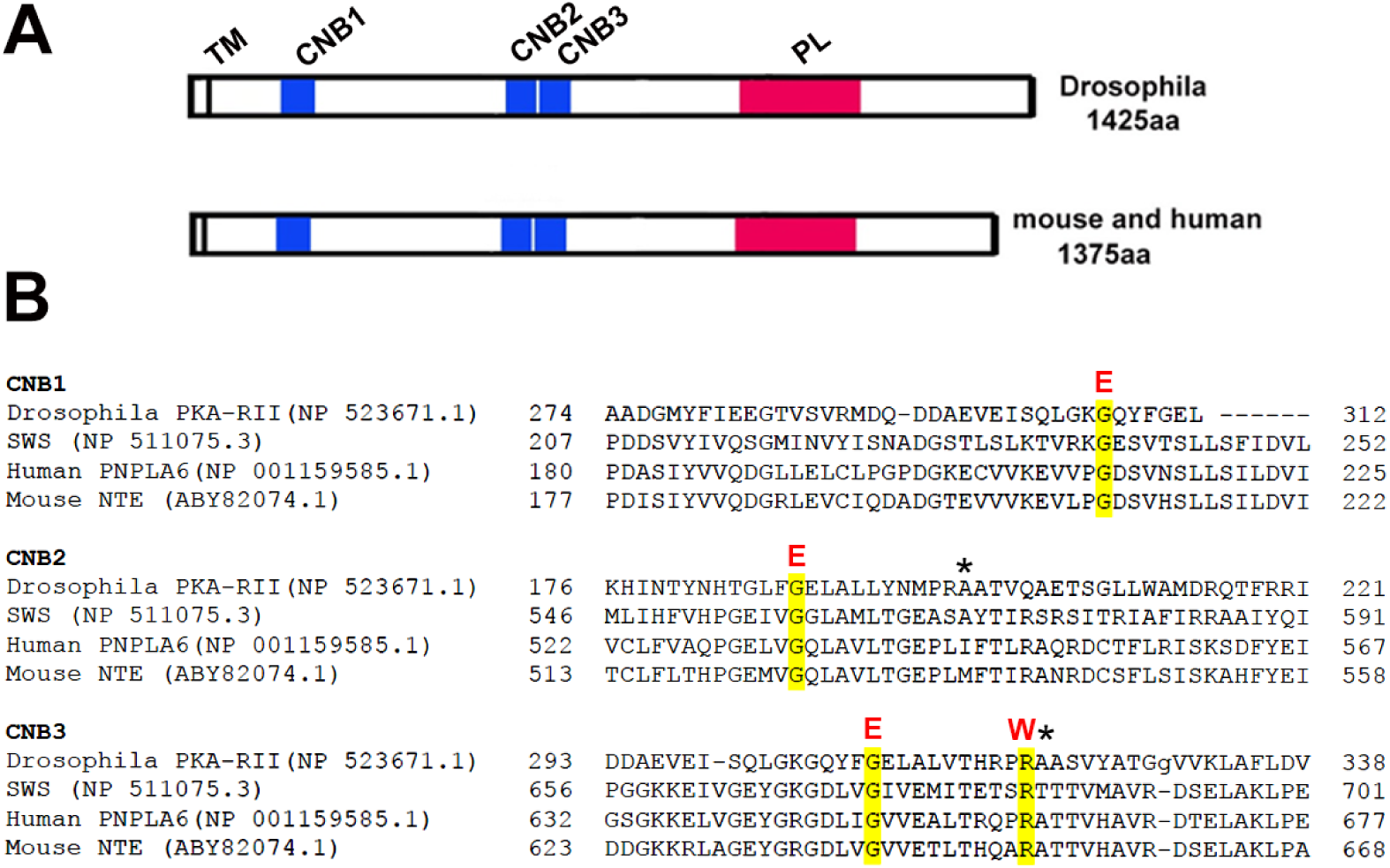
A) Schematic of *Drosophila* and mammalian SWS/PNPLA6. The cyclic nucleotide binding sites (CNBs) are shown in blue, the phospholipase domain (PL) in red, and the transmembrane domain (TM) is indicated by a vertical stripe. B) Sequence comparison of the CNB regions from *Drosophila*, mouse, and human SWS/PNPLA6 compared to the *Drosophila* regulatory PKA subunit RII. The sites mutated in the generated constructs are highlighted. The alanines indicated by asterisks are generally found in cAMP binding sites.

To determine the functional consequences of the mutations in the CNBs, we investigated whether they can revert the phototaxis phenotype that occurs when knocking down *sws* in neurons. As shown in figure 2A and B, the knockdown of *sws* in neurons via *elav*-GAL4 resulted in a significant reduction in phototaxis in 30d old female and male flies (red bars) compared to age-matched wild type Canton S (CS) and the *elav*-GAL4 driver crossed to CS (grey). Expressing SWS without a mutation in the *sws* knock-down flies rescued the phototaxis performance to the levels of the controls in females and males (Fig. 2A, B, green). Using the sws construct with the mutation only in CNB2 (SWS^G558E^) also restored normal behavior but only in females (Fig. 2A). In contrast, females expressing the SWS construct with mutations in all three CNBs (SWS^ΔCNB^) were not significantly different from the *sws* knockdown and were significantly worse than the controls and the rescue with the wildtype sws construct. In males, both mutant constructs were incapable of rescuing the phototaxis phenotype and they were significantly worse than controls and the wildtype rescue (Fig. 2B).

**Fig. 2.**
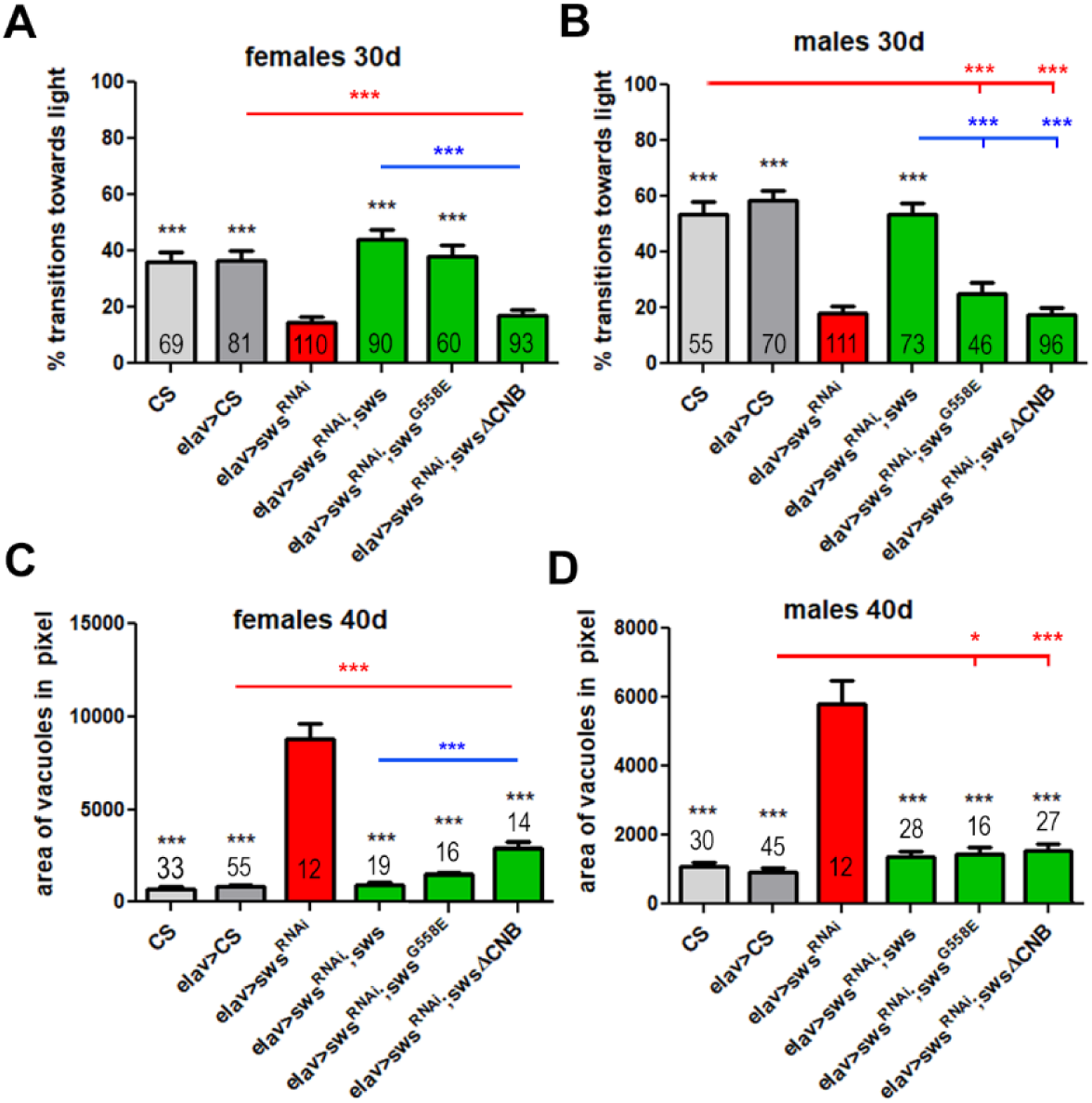
Mutating the CNB sites affects the ability of SWS to restore the wildtype function in the neuronal sws knockdown (*elav*>sws^RNAi^). A, B) When all three CNBs are mutated (SWS^ΔCNB^), the construct cannot restore phototaxis behavior while wildtype sws can. Mutating only CNB2 (SWS^G558E^) retains the function in females (A) but not in males (B). C, D) Expresssion of all constructs improves neurodegeneration but SWS^ΔCNB^ is less efficient than SWS. n is indicated in or above the bars. Error bars indicate SEMs. Comparisons were done with One-way ANOVA with a Dunnett’s post-test to the knockdown control (black asterisks), to heterozygous *elav*-GAL4 carrying flies (red asterisks) and to the rescue with wildtype sws (blue asterisks). *p<0.05, **p<0.01, ***p<0.001.

Next, we tested whether the constructs rescue the neurodegeneration occurring in the elav>sws^RNAi^ knockdown flies. As shown in figure 2C and D, knocking down *sws* caused robust degeneration, measured as the area of spongiform lesions in brain sections of 40d old females and males and this was reduced by expressing the wildtype construct (Fig. 2C, D). In contrast to the phototaxis, both SWS^G558E^ and SWS^ΔCNB^ did significantly suppressed the neurodegeneration. However, in females the rescue with SWS^ΔCNB^ was not as efficient as with wildtype SWS (Fig. 2C) and the degeneration was also still significantly worse than in the *elav*-GAL4 controls. In contrast, SWS^G558E^ was not significantly different from SWS in its rescue ability but it did not quite reduce the degeneration to the levels in the *elav*-GAL4 control in males. Together this shows that deleting all three CNB sites prevents the ability of SWS to rescue phototaxis behavior, although it still partially rescues the degeneration.

### 3.2. Overexpressing SWS in neurons causes reduced phototaxis which is prevented by mutations in the CNBs

We previously showed that overexpression of SWS in neurons also reduced phototaxis responses (Wentzell *et al*., 2014). To determine whether this requires the function of the CNB sites, we induced the mutant sws constructs with *elav*-GAL4 in the wildtype background. Testing females when 30 days old, we found a significant reduction in performance with the wildtype sws construct but both mutant constructs had no effect (Fig. 3A). In males, both SWS and SWS^G558E^ reduced the phototaxis while SWS^ΔCNB^ did not (Fig. 3B), confirming that the CNB sites are also required to induce the overexpression phototaxis phenotype. Measuring degeneration in the SWS overexpression flies did not reveal an effect of any of the constructs (Fig. 3C, D).

**Fig. 3.**
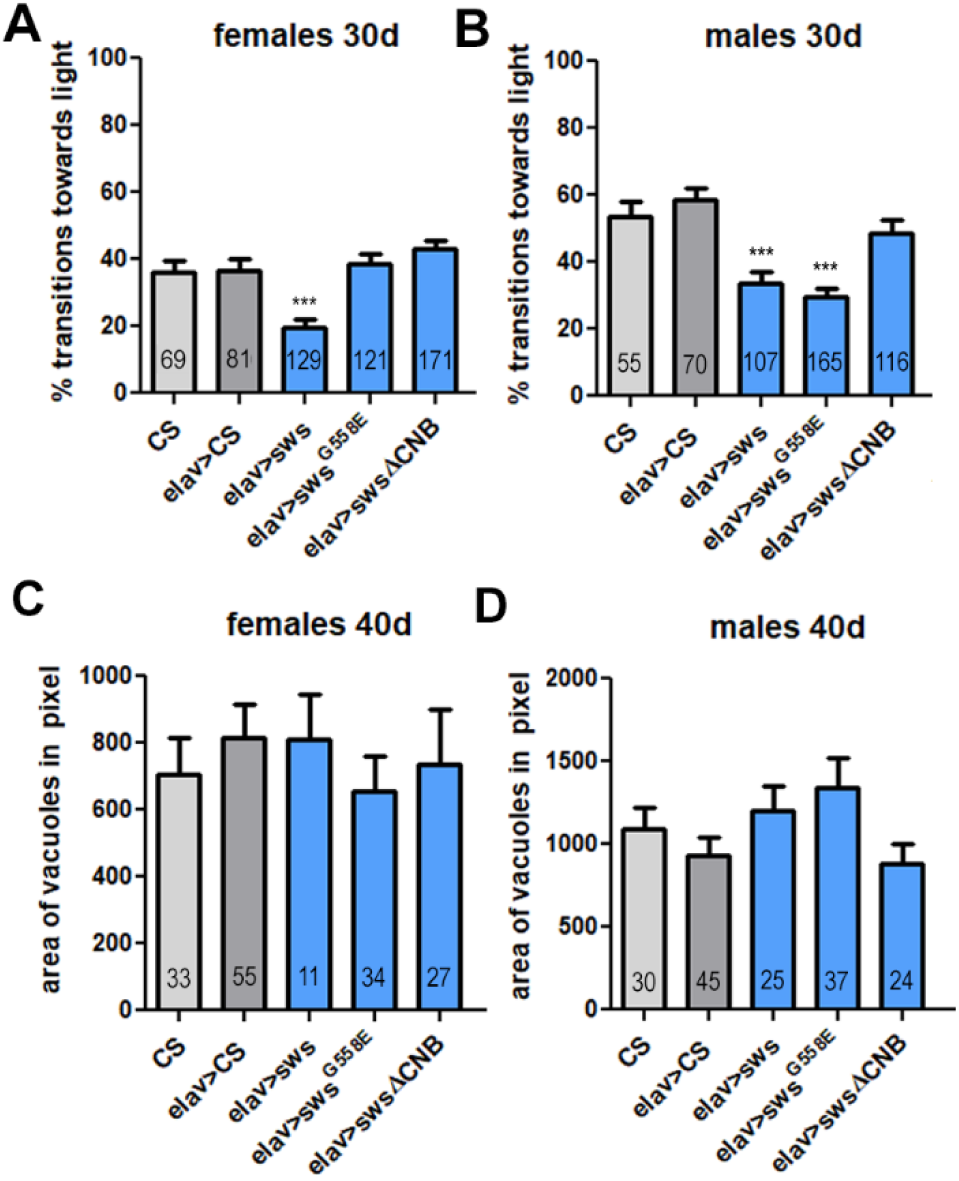
Wildtype SWS and SWS^G558E^ reduce phototaxis when expressed pan-neuronally in the wildtype background. Whereas only expression of wildtype SWS reduces phototaxis in females (A), both SWS and SWS^G558E^ reduce in males (B). C, D) None of the constructs induced degeneration when overexpressed in neurons. n is indicated in the bars. Error bars indicate SEMs. Comparisons were done with One-way ANOVA with a Dunnett’s post-test to heterozygous elav-GAL4 carrying flies. ***p<0.001.

### 3.3 The mutant SWS proteins retain a partial function when tested in the glial sws knockdown

SWS is expressed in neurons and glia and both cell types require SWS cell-autonomously (Muhlig-Versen *et al*., 2005). Specifically knocking down SWS in glia via *Gli*-GAL4 also resulted in reduced phototaxis and degeneration which was most prominent in the lamina cortex (Dutta *et al*., 2016). As seen in figure 4A and B, the phototaxis phenotype is already detectable in 14d old flies and as expected, co-expressing wildtype *sws* improved the phototaxis. However, this did not reach the performance level of the control flies (heterozygous *Gli*-GAL4). Co-expression of the mutant constructs also increased the performance but this was only significant with SWS^ΔCNB^ in females (Fig. 4A). In contrast to the experiments with the neuronal knockdown, there was no significant difference between the mutant sws constructs and the wildtype construct. Addressing the degeneration in the lamina cortex, again co-expression of wildtype SWS reduced the degeneration in 20d old flies but did not completely prevent it (Fig. 4C, D; controls do not show any degeneration). SWS^G558E^ also reduced the degeneration in both sexes whereas SWS^ΔCNB^ only reduced it in males. In females, both mutant constructs were less efficient than the wildtype construct whereas in males this was only the case for SWS^ΔCNB^.

**Fig. 4.**
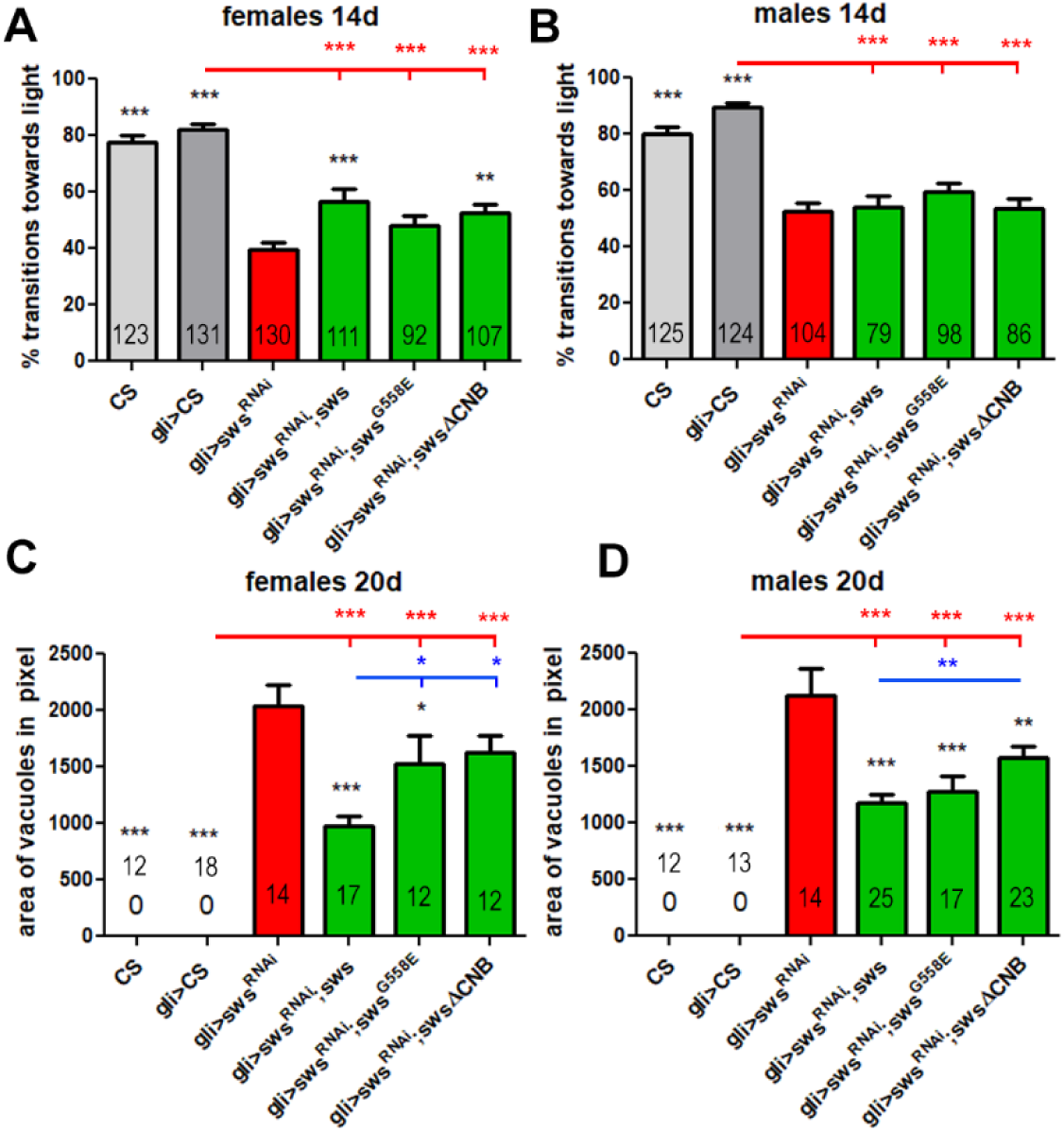
Expression of the SWS constructs had only small effects on the glial sws knockdown but can improve the degeneration in the optic lobes of these flies. A, B) Unlike the homologous experiment with *elav*-GAL4, none of the constructs fully restores SWS function in the phototaxis behavior of the glial knockdown (red asterisks compared to *gli*>CS). However, SWS and SWS^ΔCNB^ partially improve phototaxis in females compared to *gli*>sws^RNAi^ (A, black asterisks). C, D) All constructs, with the exception SWS^ΔCNB^ in females, reduce the degeneration compared to the glial knockdown (black asterisk). However, SWS^ΔCNB^ is significantly less efficient than SWS (blue asterisks). n is indicated in or above the bars. Error bars indicate SEMs. CS and heterozygous *gli*-GAL4 carrying flies show no degeneration (indicated by 0 in C and D). *p<0.05, **p<0.01,***p<0.001.

### 3.4. Overexpression of SWS and SWS^G558E^ in glia also decreases phototaxis while SWS^ΔCNB^ does not

Similar to what we observed after neuronal overexpression of SWS, we found that overexpression of the wildtype sws construct induced a phototaxis phenotype in 14d old males and females (Fig. 5A, B). This was also the case when expressing SWS^G558E^ although in males this was only significant when compared to the *Gli*-GAL4 control and not to the CS control. SWS^ΔCNB^ did not have an effect, again showing that the CNBs are required for the overexpression phenotype. None of the sws constructs caused degeneration in the lamina cortex in 14d old flies. When performing head sections of 20d old flies and measuring the area of vacuoles in all areas again none of the constructs caused increased degeneration in females compared to controls. In males, expression of any of the constructs resulted in significantly more degeneration when compared to the *Gli*-GAL4 control but not compared to CS. Together the overexpression experiments show that expression of additional SWS in either neurons or glia reduces the performance in the phototaxis assay and that this requires intact CNB sites. In contrast, even an excess of wildtype SWS expression in neurons or glia does not induce degeneration (at least when compared to both control lines).

**Fig. 5.**
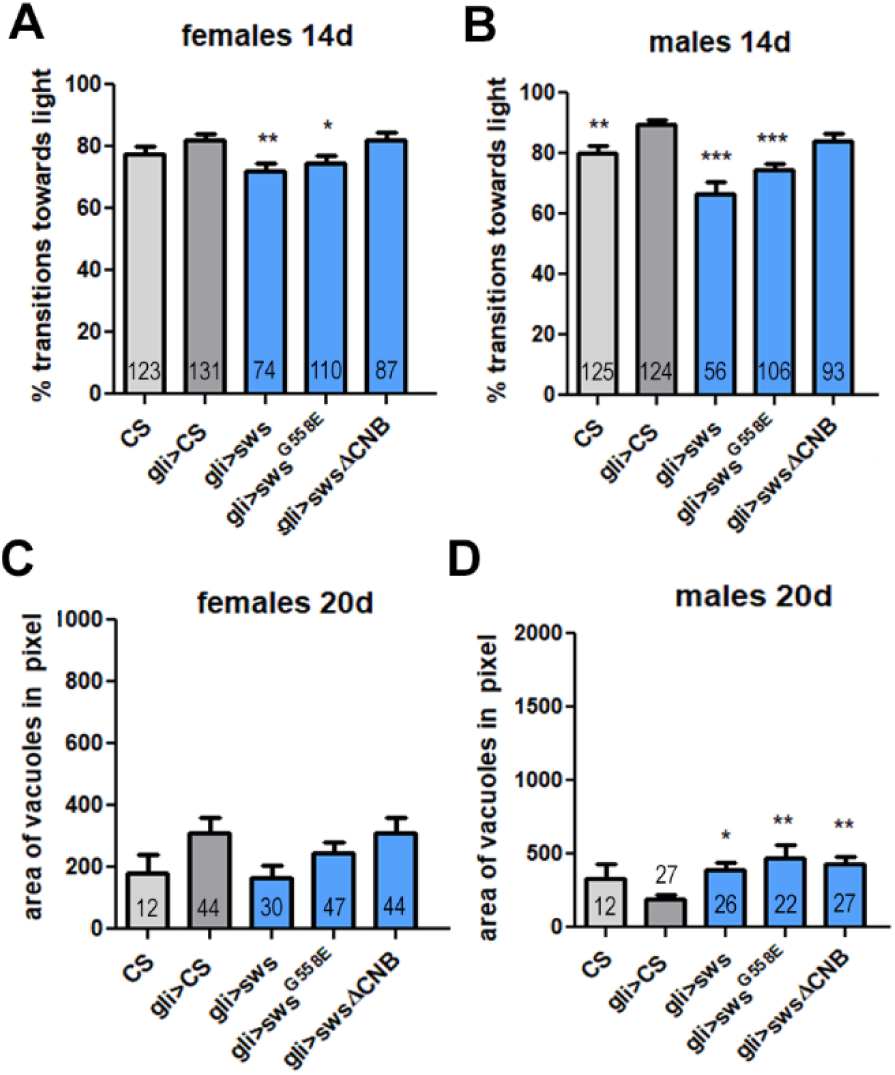
Wildtype SWS and SWS^G558E^ reduce phototaxis when expressed with gli-GAL4 in the wildtype background (A, B). C, D) None of the constructs induces degeneration in females but they do in males when compared to heterozygous gli-GAL4 flies (there is no difference to CS). n is indicated in or above the bars. Error bars indicate SEMs. Comparisons were done with One-way ANOVA with a Dunnett’s post-test to heterozygous elav-GAL4 carrying flies. *p<0.05, **p<0.01, ***p<0.001.

### 3.5. Cyclic nucleotides induce a release of PKA-C3 into the cytoplasm

We previously showed that SWS can act as a non-canonical PKA regulatory subunit, binding to the PKA-C3 catalytic subunit and tethering it to the membrane (Bettencourt da Cruz *et al*., 2008). Consequently, the loss of SWS results in less membrane bound PKA-C3, increased cytoplasmic PKA-C3 and increased PKA activity (Bettencourt da Cruz *et al*., 2008). Canonical regulatory subunits of PKA release the catalytic subunit after binding of cAMP (Taylor *et al*., 2005). However, it has also been shown that cGMP can activate PKA at least in some contexts (Sausbier *et al*., 2000; Worner *et al*., 2007). To determine whether the release of PKA-C3 from SWS is regulated by cyclic nucleotides, we determined the localization of PKA-C3 when adding cyclic nucleotides. For this, we used flies that expressed PKA-C3 with a C-terminal Myc-tag pan-neuronally with *Appl*-GAL4. Head extracts made from these flies were then incubated with cAMP, cGMP, or both before preparing cytosolic and membrane fractions and Western blots. We previously showed that PKA-C3, which is predicted as a 65kD protein, is detectable as an approximately 73kD band in Western blots in membrane fractions with an additional smaller form of about 60kD present in the cytoplasmic fraction (Bettencourt da Cruz *et al*., 2008). As shown in figure 6, we also detected bands of these sizes in the head extracts of PKA-C3-Myc expressing flies (lanes 2 and 3). When the extracts were incubated with either cAMP or cGMP or both, the membranous form was decreased, whereas the shorter cytosolic form was increased compared to the extracts without adding cyclic nucleotides. That both cyclic nucleotides induce an increase of the cytoplasmic form suggest that both, cAMP and cGMP can bind to SWS and result in the release of PKA-C3.

**Fig. 6.**
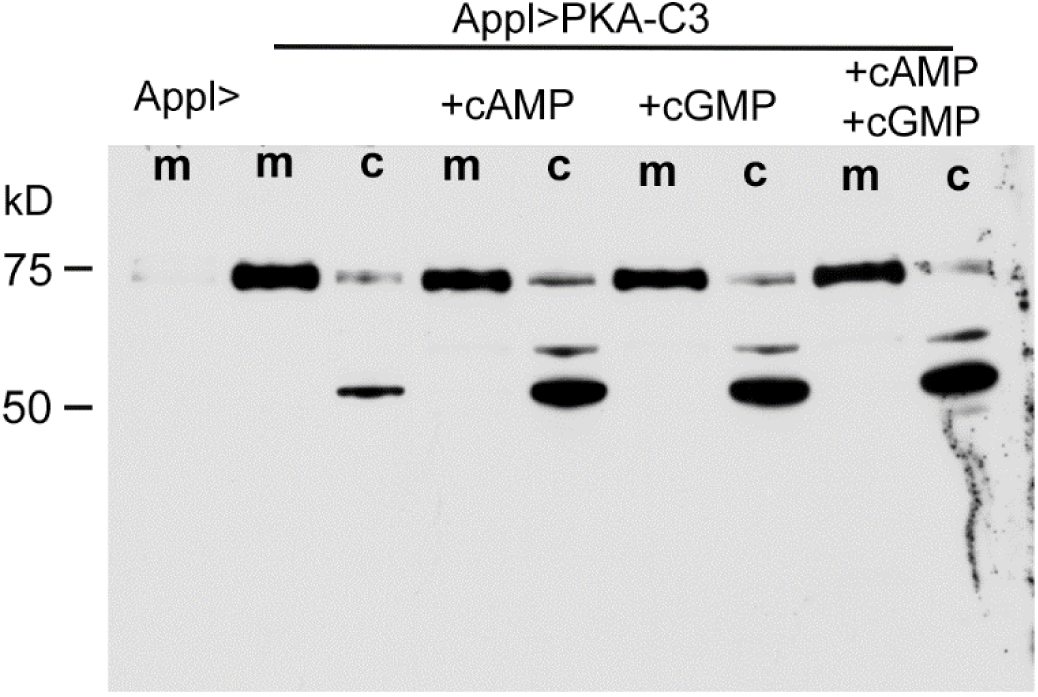
cAMP and cGMP can release PKA-C3. Adding cAMP, cGMP or both to head extracts of PKA-C3-Myc expressing flies results in the increased release of the smaller PKA-C3 form into the cytoplasm. m=membrane fraction, c=cytosolic fraction.

### 3.6. Mutations in the CNBs impact the phospholipase function

PNPLA6/NTE activity has originally been measured by using phenyl valerate as a substrate in an invitro assay (Johnson, 1977; Xie *et al*., 2003; Richardson *et al*., 2013). Using this assay, we previously found that the SWS^R133A^ mutation, that prevents the binding of SWS to the PKA-C3 regulatory subunit, does increases the catalytic activity of SWS when expressed pan-neuronally (Bettencourt da Cruz *et al*., 2008). As mentioned in the introduction, we also found that human PNPLA6 with mutations in the CNB region, cannot restore the increased levels of PC and LPC in *sws* mutant flies, although wildtype human PNPLA can at least restore the LPC levels (Sunderhaus *et al*., 2019). This suggests that binding of cyclic nucleotides and/or the release of PKA-C3 can affect the phospholipase function of SWS. To address this, we performed a lipidomic study using heads of flies expressing the sws constructs pan-neuronally (Fig. S4). As described previously (Muhlig-Versen *et al*., 2005), we found a decrease in PCs, suggesting it lost its phospholipase function when expressing wildtype SWS, although this did not reach significance (Fig. 7A). However, the decrease was significant when expressing SWS^G558E^. In contrast, SWS^ΔCNB^ significantly increased PC levels. When analyzing the levels of saturated versus unsaturated fatty acids, we found that a decrease in SWS^G558E^ and an increase in SWS^ΔCNB^ was detectable in saturated, monounsaturated, and polyunsaturated PCs (Fig. 8A). Measuring the levels of LPC species, SWS expression reduced the levels of LPC but neither mutation had a significant effect (Fig. 7B). The decrease was specifically detected in monounsaturated LPCs (Fig. 8B). Analyzing other lipids for significant changes, we found that phosphatidylserine (PS) levels were reduced by wildtype SWS and SWS^G558E^ but they were increased by SWS^ΔCNB^ (Fig. 7C). Altered levels were detected in saturated as well as unsaturated fatty acids (Fig. 8C). In addition, wildtype SWS and SWS^G558E^ decreased ceramide-phosphoethanolamine (Cer-PE) whereas SWS^ΔCNB^ increased it (Fig. 7D). Due to the role of SWS in hydrolyzing PC and LPC we expected a decrease in PC and/or LPC when overexpressing SWS and we also detected a decrease in PC when expressing additional SWS^G558E^. This suggests that SWS^G558E^mostly retains the phospholipase function of wildtype SWS. In contrast, SWS^ΔCNB^ increased PC indicating that the mutation in all three binding sites prevents the catalytic activity and even impairs the phospholipase function of the wildtype SWS that is still present in these flies.

**Fig. 7.**
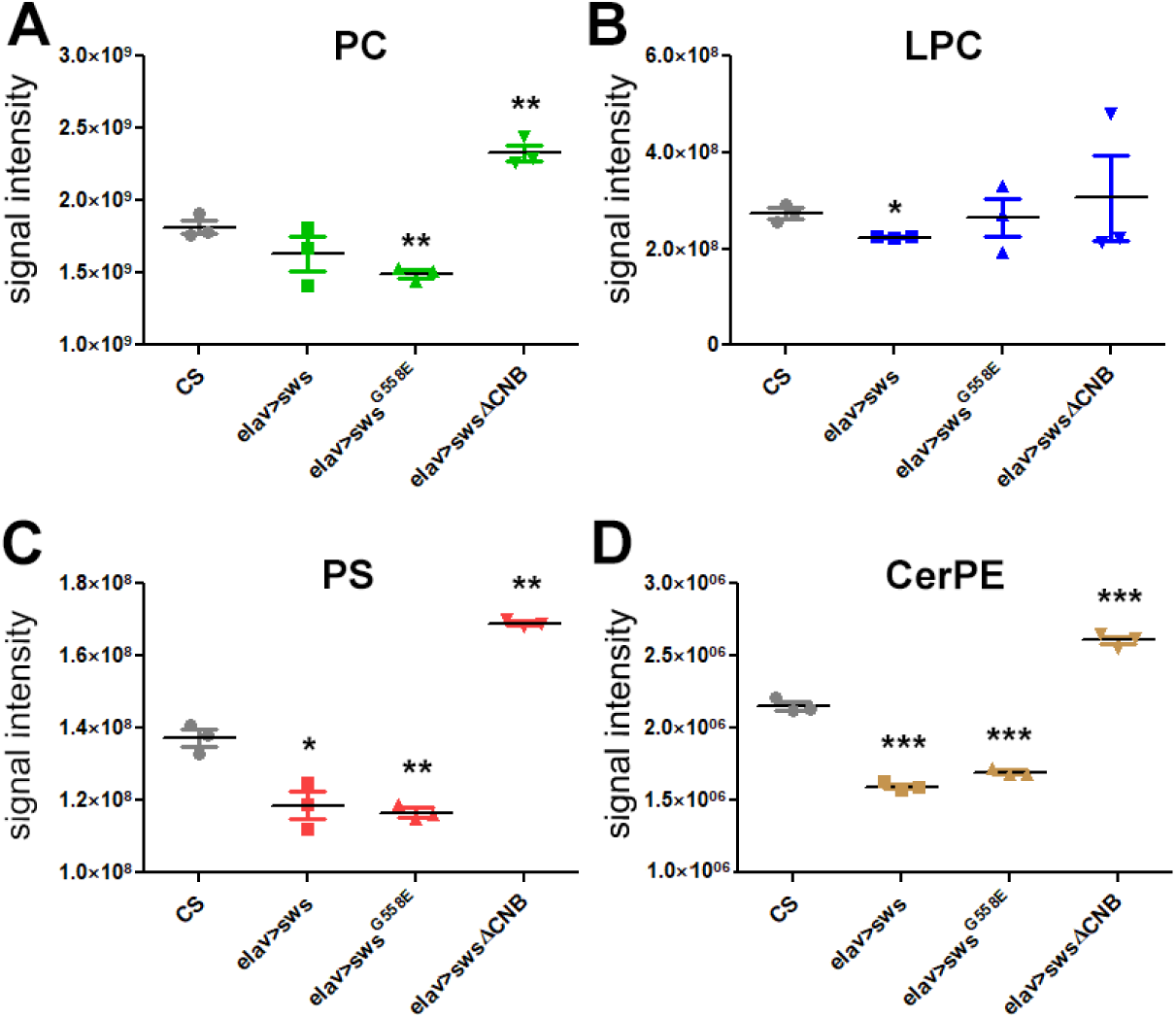
Mutations in the CNBs affect the phospholipase function of SWS. A) While pan-neuronal expression of both, SWS and SWS^G558E^ decrease phosphatidylcholine (PC) this only reached significance for SWS^G558E^. SWS^ΔCNB^ increases PC. B) Lysophosphatidylcholine (LPC) levels are only affected by SWS, decreasing them. C) Significant changes are also detected for the levels of phosphatidylserine (PS) which are decreased by SWS and SWS^G558E^ but increased by SWS^ΔCNB^. D) A decrease by SWS and SWS^G558E^ and increased by SWS^ΔCNB^ is also detected in the levels of ceramide-phosphoethanolamine (CerPE). Statistics are done using Welch’s t-test to compare to CS. n=3 for each individual lipid. *p<0.05, **p<0.01, ***p<0.001.

**Fig. 8.**
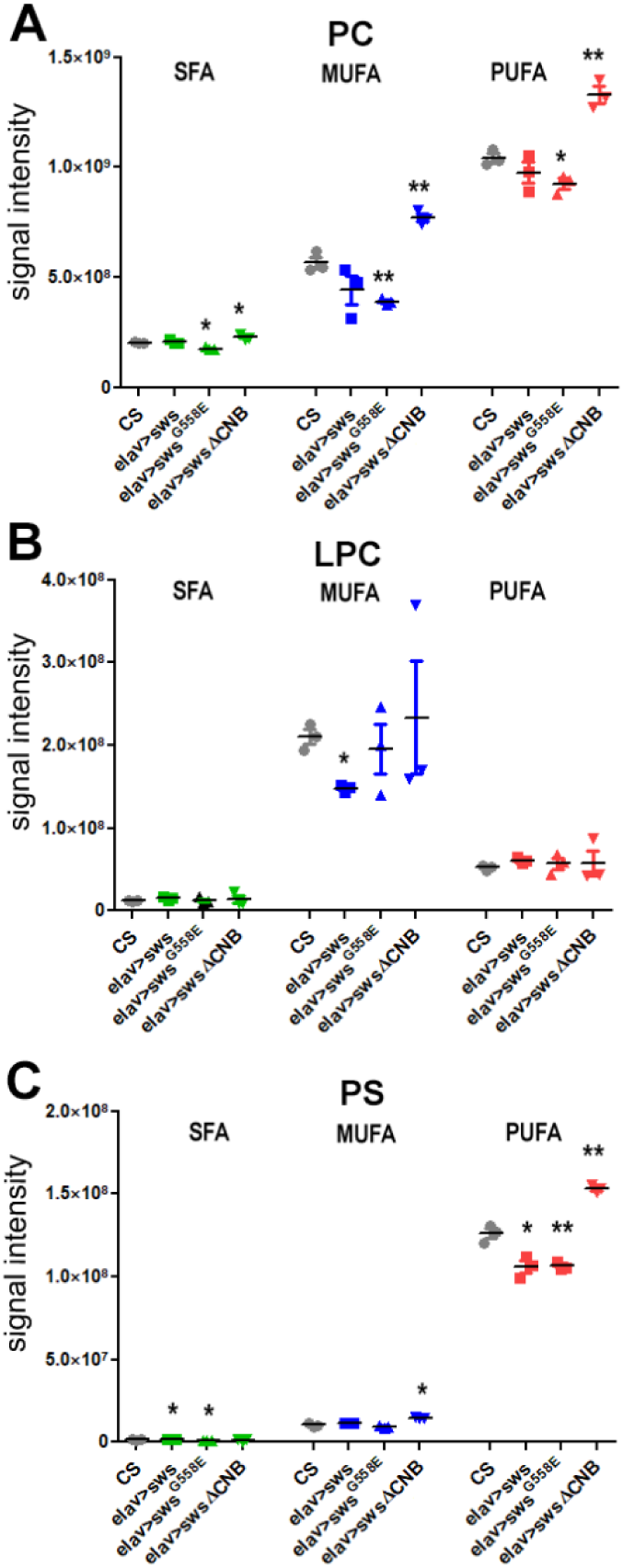
Mutations in the CNBs affect the levels of saturated (SFA), monounsaturated (MUFA) and polyunsaturated (PUFA) PC. B) In the case of LPC, only MUFAs are reduced by SWS expression. C) SWS and SWS^G558E^ affect SFA levels of PS while SWS^ΔCNB^ increases MUFAs. PUFAs are affected by all SWS constructs. Statistics are done using Welch’s t-test to compare to CS. n=3 for each individual lipid. *p<0.05, **p<0.01.

## 4. Discussion

*Drosophila* SWS and its human orthologue PNPLA6 share a conserved phospholipase domain but also three regions that show homology to cyclic nucleotide binding sites (CNBs).

Mutations in PNPLA6 cause a variety of syndromes and some of these mutations are in the CNB regions. This includes PNPLA6^L524P^, PNPLA6^G578W^, and PNPLA6^T629R^ and although they are all affecting amino acids near the CNB sites, they cause different syndromes. PNPLA6^L524P^ was identified in patients with Oliver McFarlane Syndrome, who show chorioretinopathy and hypothyroidism (Hufnagel *et al*., 2015; Kmoch *et al*., 2015; Synofzik *et al*., 2015). PNPLA6G578W causes Boucher-Neuhäuser Syndrome and these patients show cerebellar atrophy, ataxia, hypogonadism, and chorioretinal dystrophy (Synofzik *et al*., 2014). In contrast, patients with the PNPLA6^T629R^ mutation exhibit signs characteristic for Gordon–Holmes syndrome, mainly hypogonadism with absent or delayed pubertal development (Kotan *et al*., 2016). Furthermore, expressing human PNPLA6 with these mutations in Drosophila lacking SWS, showed that they cannot revert the locomotion phenotype whereas wildtype PNPLA6 can (Sunderhaus *et al*., 2019). This suggested that the CNBs and the binding of cyclic nucleotides play an important role for the function of PNPLA6. To address this, we generated two SWS constructs that carried mutations in crucial residues in the CNB binding sites with SWS^G558E^ only affecting CNB2 and SWS^ΔCNB^ affecting all three binding sites. Expressing these constructs in the pan-neuronal knockdown of sws showed that SWS^ΔCNB^ could neither prevent the phototaxis defects nor the degenerative phenotype in the knockdown flies, whereas wildtype SWS could. Although the degeneration was reduced, SWS^ΔCNB^ expressing flies still had significantly more degeneration than control flies and, in females, SWS expressing flies. We also investigated a rescue function of the SWS constructs in the glial knockdown of sws but due to much weaker effects of the wildtype SWS, the results from the phototaxis experiments with the mutant constructs were less informative. However, when analyzing the degeneration in the optic system of these flies, SWS^ΔCNB^ could reduce the degeneration but was not as efficient as SWS. In general, SWS^G558E^ showed a better rescue ability than SWS^ΔCNB^, restoring the phototaxis phenotype in females and the degenerative phenotype in both sexes in the neuronal knockdown as efficiently as wildtype SWS. With the exception of the degeneration in female flies, SWS^G558E^ was also not significantly different from wildtype SWS in the glial knockdown. SWS^G558E^, like wildtype SWS, also induced phototaxis defects in the wildtype background, further supporting the conclusion that it mostly maintained the normal function whereas SWS^ΔCNB^ did not. When comparing the effects of the mutant constructs on suppressing the phototaxis defects versus the neurodegeneration, we found that that even SWS^ΔCNB^ was quite effective in reducing the degeneration while not reducing the phototaxis. This suggests that the binding of cyclic nucleotides to SWS is of greater importance for the behavior than the integrity of the brain.

We previously showed that SWS binds to PKA-C3 and that preventing this interaction releases PKA-C3 into the cytoplasm. Using PKA-C3 localization we found that cAMP as well as cGMP induce the release of PKA-C3, further supporting that SWS acts as a non-canonical PKA regulatory subunit that releases the catalytic subunit after binding cAMP or cGMP. More surprising was the effect of the mutations in the CNBs on lipid levels. As described in the introduction, PNPLA6 hydrolyzes PC and LPC (Lush et al. 1998, van Tienhoven et al. 2002, Quistad et al. 2003)(Ono *et al*., 2025) and loss of SWS results in increased PC and LPC levels (Muhlig-Versen et al. 2005, Kmoch et al. 2015). Consequently, overexpressing SWS should result in a decrease of PC and LPC and indeed, we did detect a decrease in LPC and PC. Whereas SWS reduced both, this only reached significance for LPC, suggesting that it has a higher affinity for LPC. This seems at least to be the case for human PNPLA6 expressed in sws mutant *Drosophila*, which restored LPC levels but not PC levels (Sunderhaus *et al*., 2019) In contrast, additional expression of SWS^G558E^ did reduce PC but without a significant reduction in LPC. This suggests that it retains the phospholipase function but might be more efficient in hydrolyzing PC than LPC. Together with its ability to rescue behavioral and degenerative phenotypes in the sws knockdown, this again shows that mutating only CNB2, is not sufficient to induce a complete loss of SWS function. In contrast, SWS^ΔCNB^ resulted in an accumulation of PC, presumably because it has lost or reduced phospholipase activity. This suggests that either CNB binding activates the phospholipase directly or alternatively the release of PKA-C3 is needed for the lipase activity. Analyzing the levels of other lipids, we found that expression of the SWS constructs also changed phosphatidylserine (PS) levels. So far effects of PNPLA6/SWS on PE have not been well described however, a lipidomic study of PNPL6 deficient human ARPE-19 cells identified an increase in PS (Ono *et al*., 2025). This is in agreement with our findings that SWS and SWS^G558E^ overexpression reduces PS and that SWS^ΔCNB^, which appears to have lost its phospholipase function, increases PE. PS can be synthesized from PC by phosphatidylserine-synthase (PSS) in mammals (Vance & Steenbergen, 2005) and this enzyme is conserved in *Drosophila* (Park *et al*., 2021). It is therefore possible that the changes in PS are due to the changes in PC. Alternatively, PNPLA6/SWS can hydrolyze PS in addition to PC and some phospholipases can cleave several phospholipids (Bonting *et al*., 1980; Pete & Exton, 1995). In contrast to PC, PS levels in the membrane are relatively low. Nevertheless, it has key functions in recruiting and activating several enzymes and in apoptosis (Leventis & Grinstein, 2010). The importance of PS has also been shown using mutants of PSS in *Drosophila* which show a reduced lifespan and climbing defects (Park *et al*., 2021). It is therefore possible that the phototaxis deficits in SWS and SWS^G558E^ overexpressing flies are due to the decreased PS levels. Lastly, we found a decrease in ceramide-phosphoethanolamine (Cer-PE) in SWS and SWS^G558E^ expressing flies while it was increased with SWS^ΔCNB^. Cer-PE is the major sphingolipid in invertebrates but can also be found in mammals, although at very low levels (Panevska *et al*., 2019). In *Drosophila* Cer-PE is produced from CDP-ethanolamine by the ceramide phosphoethanolamine synthase (CPES) and cpes mutant flies have seizures, are short-lived, and show incomplete neuronal ensheathment by glia (Kunduri *et al*., 2018). They also have high levels of LPC, providing a connection between Cer-PE and the phosphatidylcholine pathway. Like PE, PC can be generated by the Kennedy pathway from CDP-choline, which is then hydrolyzed by PNPLA6 (Fagone & Jackowski, 2013; Liu & Hufnagel, 2023). Interestingly, like PNPLA6, enzymes in the Kennedy pathway that generate PE, have also been shown to cause motor neuron diseases when mutated (Rickman *et al*., 2019). However, whether the Kennedy pathway plays a role in the effects of PNPLA6/SWS on Cer-PE levels remains to be determined. In general, we detected the most prominent effects on lipids in SWS^ΔCNB^ with a significant increase in glycerophospholipids, including PC, PE, PI, PG, and PS (Table S1) whereas SWS and SWS^G558E^ either had no significant effect or decreased them.

## Supporting information

lipidomics table

## Acknowledgement

This work was supported by NIH grants (NS047663) to D.K. and (1S10OD032323-01) to L.M.

