## supplemental figures for "The cyclic nucleotide binding sites of Swiss-Cheese, the *Drosophila* orthologue of human PNPLA6, are required for its catalytic function"

### Supplementary Figures

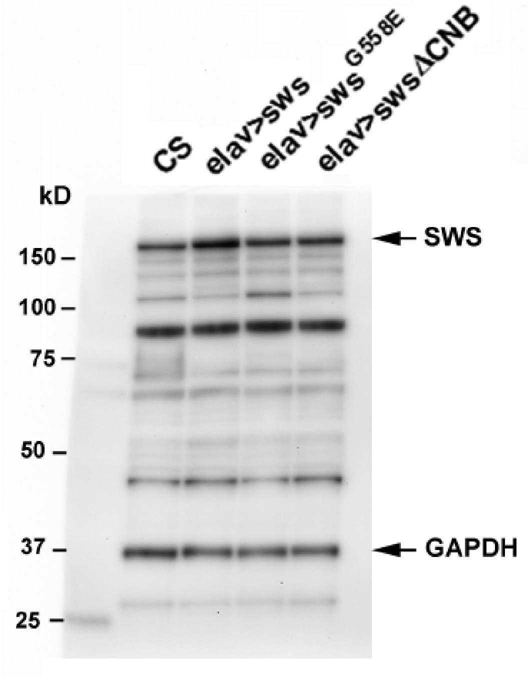


**Supplementary figure 1) Western blot showing expression of the mutant sws constructs. Their expression levels are lower when compared to sws. The loading control using anti-GAPDH shows a higher load in CS.**

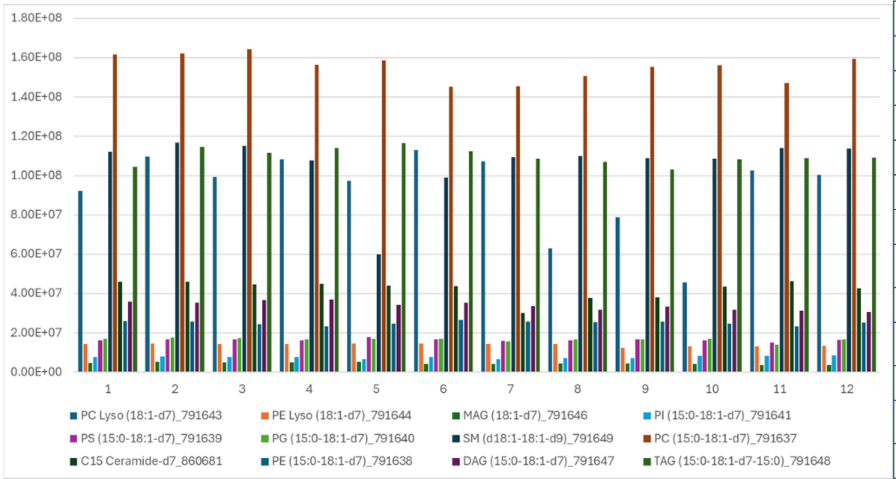


| EquiSPLASH Standard Lipid | %RSD |
|---|---|
| PC Lyso (18:1-d7) | 22% |
| PE Lyso (18:1-d7) | 6% |
| MAG (18:1-d7) | 12% |
| PI (15:0-18:1-d7) | 8% |
| PS (15:0-18:1-d7) | 4% |
| PG (15:0-18:1-d7) | 6% |
| SM (d18:1-18:1-d9) | 14% |
| PC (15:0-18:1-d7) | 4% |
| C15 Ceramide-d7 | 11% |
| PE (15:0-18:1-d7) | 4% |
| DAG (15:0-18:1-d7) | 6% |
| TAG (15:0-18:1-d7-15:0) | 4% |

**Supplementary figure 2) Reproducibility of internal standard (EqSPLASH LIPIDOMIX, Avanti Research) measured in positive ionization mode at 1 µg/mL. Bar graph shows the peak area for each individual measurement in order of injection. 1-3 CS, 4-6 elav>sws, 7-9 elav>sws$^{G558E}$, 10-12 elav>sws$^{\Delta CNB}$**

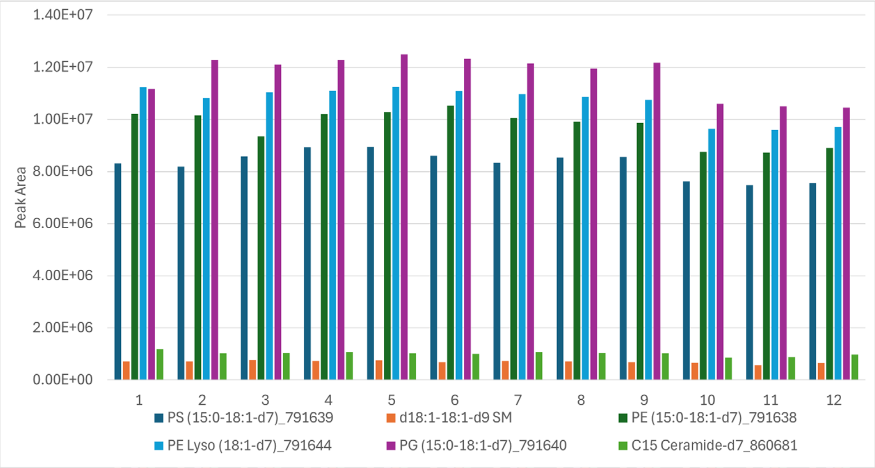


| EquiSPLASH Standard Lipid | %RSD |
|---|---|
| PS (15:0-18:1-d7) | 7% |
| SM d18:1-18:1-d9 | 9% |
| PE (15:0-18:1-d7) | 7% |
| PE Lyso (18:1-d7) | 6% |
| PG (15:0-18:1-d7) | 7% |
| C15 Ceramide-d7 | 9% |

**Supplementary figure 3) Reproducibility of internal standard (EqSPLASH LIPIDOMIX, Avanti Research) measured in negative ionization mode at 1 µg/mL. Bar graph shows the peak area for each individual measurement in order of injection. 1-3 CS, 4-6 elav>sws, 7-9 elav>sws$^{G558E}$, 10-12 elav>sws$^{\Delta CNB}$**

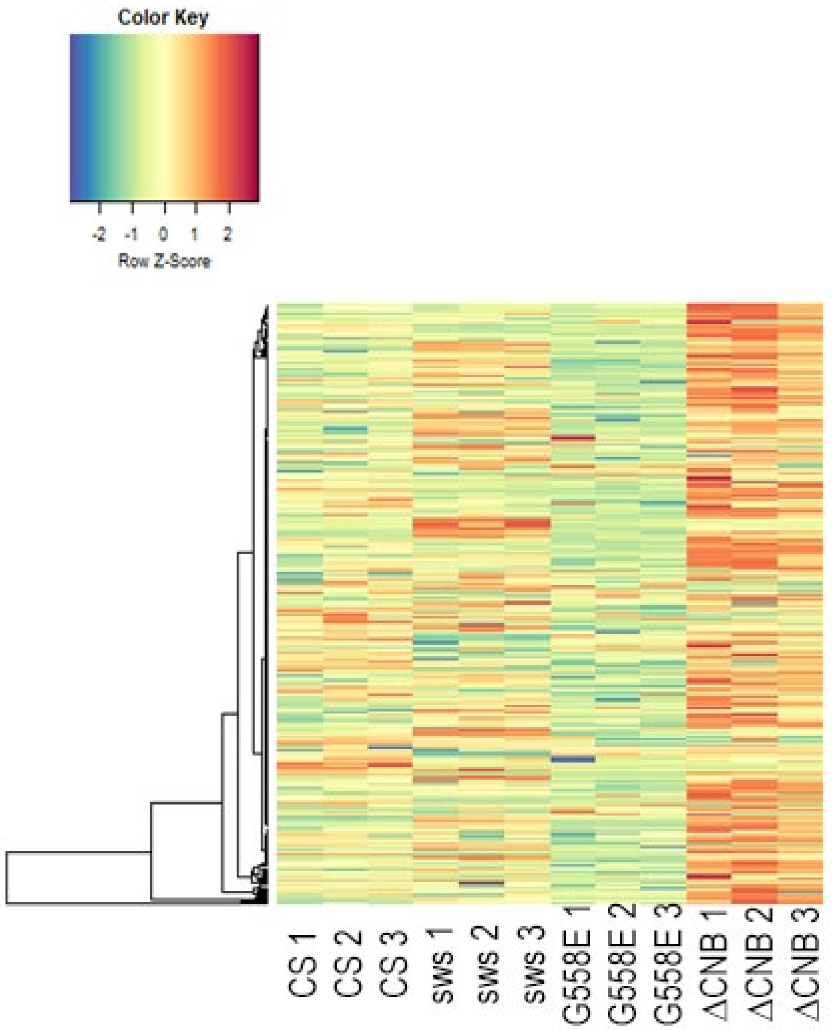


**Supplementary figure 4) Heat map showing the differences in lipids when expressing SWS, $SWS^{G558E}$, and $sws^{\Delta CNB}$. Three independent experiments were done for each genotype. Z-score (# of standard deviations from row mean) is indicated.**
